# Deep learning image analysis for filamentous fungi taxonomic classification: Dealing with small data sets with class imbalance and hierarchical grouping

**DOI:** 10.1101/2023.06.19.545596

**Authors:** Stefan Stiller, Juan F. Dueñas, Stefan Hempel, Matthias C. Rillig, Masahiro Ryo

## Abstract

Deep learning applications in taxonomic classification for animals and plants from images have become popular, while those for microorganisms are still lagging behind. Our study investigated the potential of deep learning for the taxonomic classification of hundreds of filamentous fungi from colony images, which is typically a task that requires specialized knowledge. We isolated soil fungi, annotated their taxonomy using standard molecular barcode techniques, and took images of the fungal colonies grown in petri dishes (n = 606). We applied a convolutional neural network with multiple training approaches and model architectures to deal with some common issues in ecological datasets: small amounts of data, class imbalance, and hierarchically structured grouping. Model performance was overall low due mainly to the relatively small data set, class imbalance, and the high morphological plasticity exhibited by fungal colonies. However, our approach indicates that morphological features such as color, patchiness, and colony extension rate, could be used for the recognition of fungal colonies at higher taxonomic ranks (i.e., phylum, class, and order). Model explanation implies that image recognition characters appear at different positions within the colony (e.g., outer or inner hyphae), depending on the taxonomic resolution. Our study suggests the potential of deep learning applications for better understanding the taxonomy and ecology of filamentous fungi amenable to axenic culturing. Meanwhile, our study also highlights some technical challenges in deep learning image analysis in ecology, highlighting that the domain of applicability of these methods needs to be carefully considered.

## Introduction

Deep learning is an AI toolbox inspired by the human brain to learn patterns from complex data. It is one of today’s most rapidly growing technical fields (Jordan and Mitchell, 2015). Deep learning has been used successfully in various scientific fields, including biology and medicine (Ching et al., 2018). In ecology, deep learning is used, for instance, for identifying species and classifying animal behavior from camera images, audio recordings, and videos (Christin et al., 2018).

Taxonomic classification with deep learning image analysis has been demonstrated successfully for many taxa, but applications have been limited to individuals with rigid morphological structure, such as plants, animals, and also fruiting bodies of fungi (Miao et al. 2019; Seeland et al. 2019; Hansen 2020; Dong et al. 2021). No study has investigated the potential of the technique to study the taxonomy of filamentous fungi that do not produce macroscopically visible reproductive structures, which constitute the large majority within the kingdom. Indeed, it is largely unknown whether filamentous fungi can be taxonomically classified based solely on the macroscopic morphological features when grown in a pure culture. While it is clear that taxonomic identification of an isolate to the level of species is impossible without comparing characters microscopically (keys mostly focus on fungal spores and spore producing structures, e.g. (Domsch, 2007)), or even at the molecular level, identification at a coarser taxonomic resolution might be feasible. Indeed, some early diverging lineages of filamentous fungi produce consistent colony morphologies in culture (e.g., Mortierellomycota, (Naranjo-Ortiz and Gabaldón, 2019)). Recent work suggests that hyphal growth speed (which has an impact on colony size in the Petri dish) and the complexity of the hyphal architecture are phylogenetically conserved, at least at the phylum level (Lehmann et al., 2020). Because colony characters such as color, shape, exudation, and sporulation are easily quantifiable morphological traits, we wondered whether annotating the coarser levels of the taxonomic hierarchy to a fungal colony is possible based on images using these attributes.

We used the appearance of ‘colonies’ of fungal individuals grown on Petri dishes, representing a higher level of morphological organization than hyphal traits, to test this idea. Because it is known that pure cultures of the same species can exhibit a staggering variation of morphology depending on the growing conditions, we employed a standardized culturing protocol to produce a large number of colony images. Our study first examined the application of deep learning models to achieve the taxonomic classification of filamentous fungi from the images of fungal colonies grown on Petri dishes. In particular, we asked (i) up to which taxonomic resolution (from phylum, class, order, family, genus, to species level) a deep learning model can keep accuracy, and (ii) from which region of the individual fungal colony the model learns the morphological characteristics for taxonomic classification.

In addition to these questions, we attempted to tackle some technical challenges common in ecological datasets. The size of a dataset is usually small (10^2^–10^3^) since experimental tasks are labor intensive, generally resulting in low model performance. The data containing missing values are not easy to impute in a meaningful way (e.g., many fungi cannot be identified to species level). The probability distribution of categories is highly imbalanced (e.g., many photo images belong to a handful of taxonomic groups). The classification property is hierarchically nested (e.g., once the species name is known, its higher taxonomic names are determined because of phylogeny). Since these data properties are pretty standard in ecological datasets, we think overcoming some of the technical challenges is valuable for exploring the potential application of deep learning image classification in ecology in a broader context.

## Methods and materials

### Dataset: FunTrait image collection

We obtained a subset of fungal culture images from a culture collection isolated from the German Biodiversity Exploratories’ very intensive research plots (VIPs, (Fischer et al., 2010)). These are located on grasslands at three different areas across Germany (nine per site, n = 27). VIPs represent a replicated gradient of grassland management intensity that ranges from near-pristine to heavily managed for agricultural purposes. A total of 500 g of soil from the upper 10 cm beneath the surface (e.g. (Vályi et al. 2015)) were collected per plot. Each sample represents a composite of 5 random soil cores across each plot. Upon collection, soil samples were transported to the Institute of Biology at Freie Universität Berlin, where they were stored in the dark at 4°C.

Using an in-house high-throughput culturing approach, we obtained fungal colonies from soil samples. All work, from dilution to colony isolation, was done under sterile conditions. Briefly, the soil was subjected to serial dilutions to hinder the recovery of highly sporulating fungi.Several morphologically contrasting fungal isolates were recovered from each diluted soil solution using potato dextrose agar (PDA, X931.2. Roth) in full and 1/10 strength, incubation at 12°C and subsequent transfer on new agar plates.

The cultures used to produce images grew for approximately 10-20 days at ambient temperature (approximately 15–20 degrees Celsius) in six cm diameter Petri dishes plated with 100% (w/v) sterilized malt extract agar (MEA, X923.1 Roth). To hinder bacterial growth, we mixed the sterilized MEA solution with a range of antibiotics (Penicillin G 24 μg/L, Chlortetracycline 48 μg/L and Streptomycin 26 μg/L) before casting.

Each isolate was then taxonomically annotated via DNA extraction and Sanger sequencing. Before sequencing, the entire extent of the ITS and LSU regions within the rDNA operon was amplified employing the ITS1 (Gardes & Bruns, 1993) and NL4 (O’Donnell, 1993) primers. The ITS variable regions were then extracted *in silico* with ITSx (Bengtsson-Palme, et al 2013). Finally, the blast+ classification algorithm (Camacho et al., 2009) was employed to query the ITS sequences of each isolate to UNITE’s reference database from February 2020 (Abarenkov et al., 2020a). Because UNITE databases employ the taxonomic framework proposed by Tedersoo et al. (2018), taxonomic identities inherited by the query sequences follow this framework. Each isolate was identified to the finest taxonomic level possible.

Because the goal of image identification was ancillary to the main objective of the fungal collection, Petri dishes were arranged in a semi-standardized fashion to obtain a group image. Images were group scans of up to 12 Petri dishes over a blue background screen (Fig. 1). Plates were distributed in a 3 x 4 matrix such that their position on the image corresponded to the position of their DNA extract on a 96 well DNA extraction plate. It allows the correspondence of an ITS sequence with an individual image of an isolate. Petri dishes were scanned with a Perfection V800 scanner (Seiko Epson Corporation, Japan) from the bottom and top. At the bottom of each plate, a handwritten mark with the isolate code was present. The images were formatted as file types .tif, .jpg or .png and in resolutions 2550 x 3509, 5100 x 7019 or 6800 x 9359. Only top view images were included. The dataset was composed of 606 fungal isolates as of 18th March 2020, when we established our initial study (Fig. 2).

**Figure 1.**
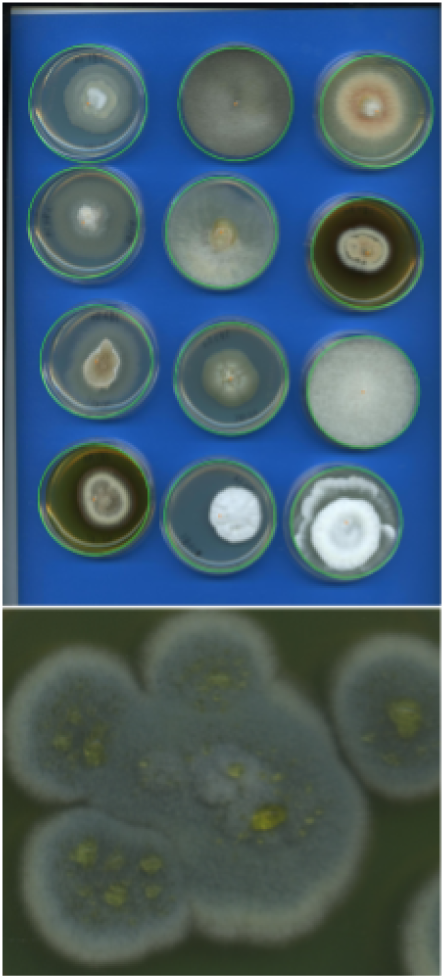
Example of a scanned image of cultures in the dataset (upper). Circles denote Petri dish object detection using the Circle Hough Transform. A zoom in on the bottom-left sample as an example of a filamentous fungus (lower).

**Figure 2.**
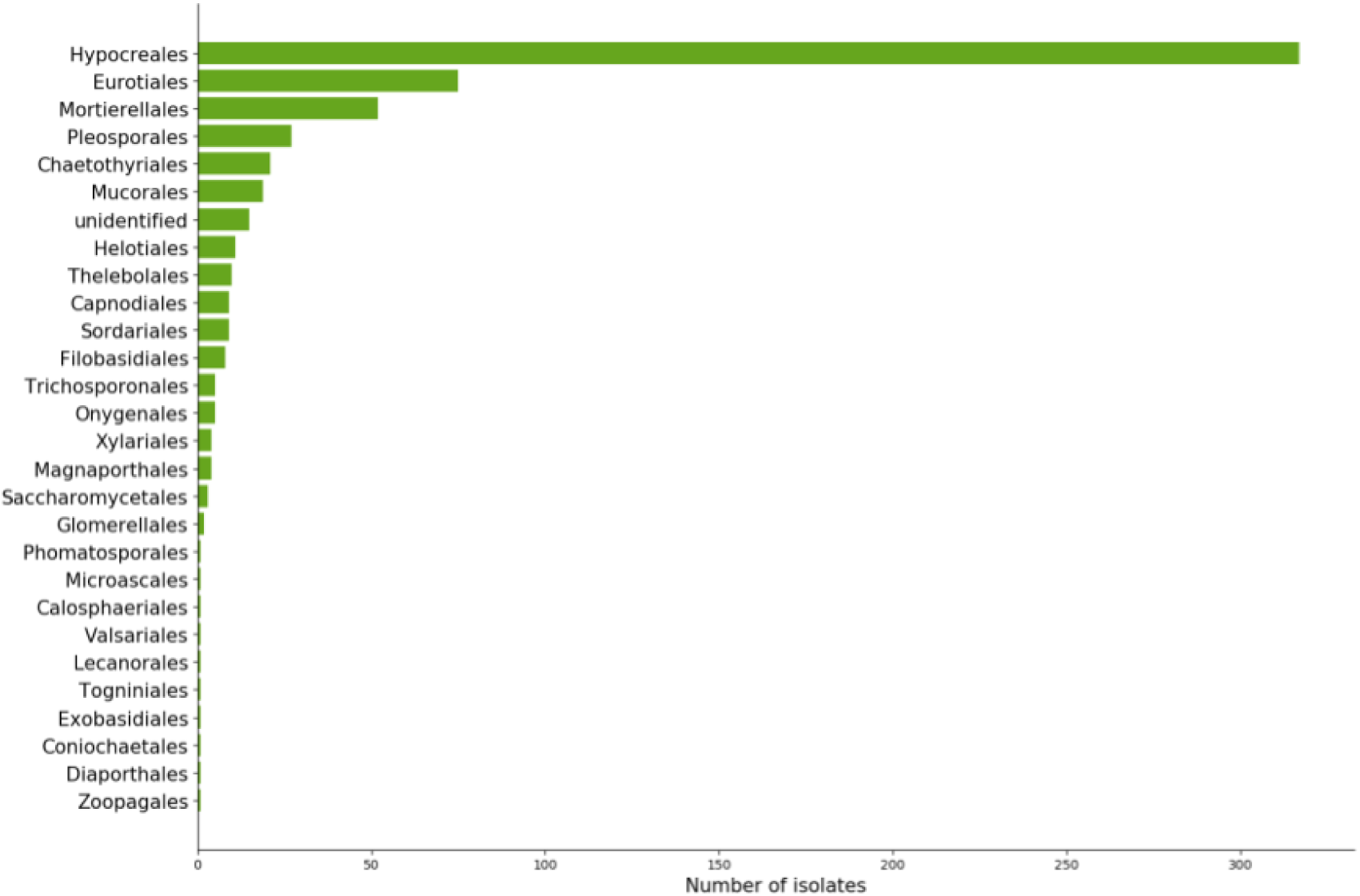
Absolute number of isolates assigned to a known order by DNA extraction and Sanger sequencing method.. Fungal isolates were extracted from soil samples, and the cultures were bred in Petri dishes. Taxonomic identification was annotated using blast+ classification algorithm, querying isolate ITS region to UNITE database.

Biological surveys inherently suffer from class imbalance, meaning that only a few taxonomic groups are highly abundant within the dataset. In the present case, molecular methods classified isolates into 5 phyla (Ascomycota, Basidiomycota, Mucoromycota, Mortierellomycota, Zoopagomycota), 11 classes, and 190 species (Fig. 3a). Because of the hierarchical organization of taxonomy, class imbalance propagates throughout the ranks, such that an imbalance occurring at a given taxonomic rank is affected by the imbalance of the preceding rank (Fig. 3b-g). Another problem of taxonomic classification by molecular methods is missing values. Missing values can occur when an isolate can be assigned to a higher taxonomic level (e.g. class) but not reliably to any of the lower taxonomic levels (order, family ect.). In this dataset, we had missing values for 11 instances for phylum, 39 for class, 40 for order, 67 for family, 119 for genus, and 319 for species. We replaced the missing values by taking the taxonomic name of the higher rank with a suffix indicating the current rank (e.g., for Ascomycota, Ascomycota_class, …, Ascomycota_species), as we were concerned that the missing values might introduce a significant bias.

**Figure 3.**
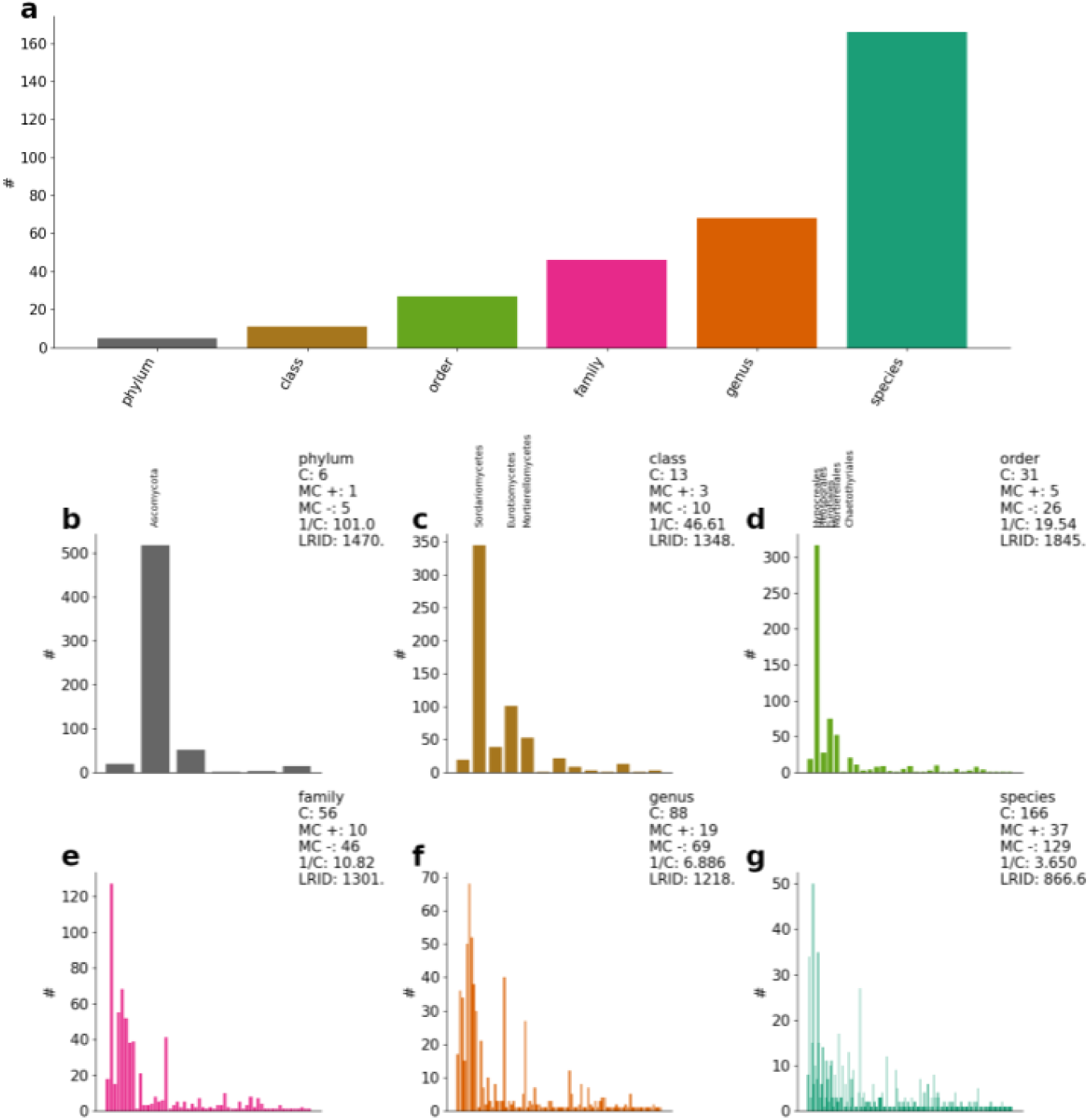
(a) Number of unique taxa/classes according to taxonomic rank for 606 samples, (b-g) the frequency distribution within each taxonomic rank. C is the number of categories; MC + is the majority category count, MC - is the minority category count. A category is regarded as a majority category if the number of samples # is higher than the average number of samples per category. LRID is the likelihood ratio imbalance degree. For phylum, class, and order levels, the majority classes are labeled for visualization.

### Image preprocessing

At first, the group image was cut, such that each sub-image contains one Petri dish scan. The images were resized to 224×224 pixels using pixel area relation for reducing the computational burden (except for augmented random oversampling to 356×356 pixels; see below). We used the Circle Hough Transform to automatically detect all Petri dishes’ positions in a single image. We used the implementation by the Open CV Project (*OpenCV 4.4.0*), which uses Hough Gradient (Petković and Lončarić, 2015) (Fig. 1). Then, each identified sample was annotated by a set of six hierarchically nested labels, each representing a taxon given a taxonomic rank, which is phylum > class > order > family > genus > species.

### Three datasets for model training

We prepared three datasets for model training to deal with class imbalance: (i) the original data as preprocessed above; (ii) naive random oversampling; and (iii) augmented random oversampling.

#### Original data (without resampling)

The images were composed of 224×224 pixels. Each image contains a single individual that is taxonomically annotated.

#### Naive random oversampling

One approach to class imbalance is to use under- and oversampling to equalize the class distributions (Batista et al., 2003). In undersampling, samples are randomly drawn from the original set down to a lower limit. Oversampling is its counterpart, where samples are randomly drawn and re-added to the setup up to a higher limit. Undersampling is not the approach of choice due to the already small size of the dataset. The problem would almost become a one-shot learning problem, which is increasingly difficult to solve. In naive random oversampling, the distribution is balanced out by increasing the sample size in the minority classes by adding slightly altered samples drawn from the minority classes while retaining the taxonomic annotation of the donor. For image alteration, the samples have their axes randomly flipped and their lighting adjusted.

#### Augmented random oversampling

Augmented random oversampling applies naive random oversampling and additionally two image augmentation techniques. The first augmentation was done at the image preprocessing stage. The high-resolution images were not resized to 224×224 pixels but 356 x 356 pixels. Then, from a 356×356 pixel image, multiple 224×224 pixel images were generated by cropping the image at random positions. It can be interpreted as zooming into a part of the image. The second augmentation was conducted on the images by applying a color jitter concerning brightness, contrast, and saturation.

## Modeling with Deep learning

### Algorithm

We applied a convolutional neural network algorithm, DenseNet-169 (Huang et al., 2017), with transfer learning to deal with the small sample size. DenseNet has some compelling technical advantages: the method is robust to the vanishing-gradient problem, strengthens feature propagation, encourages feature reuse, and substantially reduces the number of parameters. We applied transfer learning (Goodfellow et al., 2016). Transfer learning is a technique that uses a model that is already pre-trained with a vast dataset for making a model efficient. Pre-trained models were taken from MXNET Gluon Model Zoo (MXNET Gluon Model Zoo. Classification — gluoncv 0.11.0 documentation).

### Model architecture

We tried three different model (classifier) architectures run with the prepared three datasets: Separate Local per-level classifiers (SL), Multi-Label classifier (ML), and Hierarchically-Chained Local per-level classifiers (HC). These differ in how to deal with the information of the hierarchically nested structure of taxonomic classification. The model architecture comparison can be regarded as our virtual experiment to explore how to deal with (or whether it is worth dealing with) the hierarchical nature of taxonomic classification using deep learning. Silla and Freitas (2011) suggest several concepts to engage hierarchical classification, and we applied three of them in this study.

SL is the simplest approach where each taxonomic rank has its own trained classifier. There are six models (e.g., one for phylum, one for class, etc.). Therefore, the information is not passed on between taxonomic ranks, such that hierarchical information content is completely lost. Each model experiences independent training and thereby develops its separate loss function. SL utilizes one softmax output unit. They operate locally for each taxonomic rank, and hence the used output units differ as the number of possible labels differs between ranks.

ML is a single, relatively complex model, which takes the entire class hierarchy as a whole into account during a single run (Silla and Freitas 2011). It is a type of so-called global classifier or big-bang approach. A global classifier comes with the advantage of having a broad scope, i.e., learning from the hierarchical structure and possibly learning from its information. The model structure might learn the hierarchical taxonomic structure on its own through the training phase. ML uses six softmax output units, one for each taxonomic rank. However, the output units are independent of each other, meaning that the model may predict a pair of species and genus groups that can never happen together.

HC is the most complex model in this study. Based on taxonomic rank, a deep neural network classifier learns to discriminate taxons. Then, the learned parameters are passed on to the hierarchically-nested successive rank classifier, making use of transfer learning. HC makes the direct use of the taxonomic hierarchy information with six hierarchically stacked classifiers C_Phylum_ < C_Class_ < C_Order_ < C_Family_ < C_Genus_ < C_Species_. Each classifier C has its nested loss function *J*_Rank_ with respect to hierarchy, *s.t. J*_Species_ = *J*_Species_(*J*_Genus_(*J*_Family_(*J*_Order_(*J*_Class_(*J*_Phylum_))))). Six HC models apply one output unit each. Those units are in the sense that the preceding level’s output directly influences this level’s units.

The model architectures with hyperparameter settings are as follows. As specified by Huang et al. (2017) DenseNet-169 utilizes rectified linear units (ReLU) (Glorot et al., 2011), Batch Normalization (Ioffe & Szegedy, 2015), and pooling (LeCun et al., 1998). We applied L2 regularization (Ng, 2004) with weight decay *wd* = 0. 01, learning rate *lr* = 0. 001, and momentum *m* = 0. 8. Parameter initialization follows the approach Glorot & Bengio(2010), Xavier initialization. We trained the models using a softmax cross-entropy loss function (Cox, 1958).

### Model training and performance evaluation

Models are trained to maximize their performances. Therefore, the selection of a performance indicator is crucial for training. We use Accuracy and Matthews correlation coefficient (MCC). Accuracy is the most widely used indicator for classification tasks. It calculates the number of correct predictions divided by the total number of predictions and accounts for only true positive and true negative. It ranges from 0 (no correct prediction) to 1 (perfect prediction). As a critical issue of this indicator, it is sensitive to class imbalance, which our study faces. Accuracy can be high even if the model does not correctly predict rare categories at all when a few major categories occupy the data.

MCC can solve the limitation of Accuracy, as suggested by Chicco & Jurman (2020) to use as an alternative of Accuracy for imbalance datasets. MCC indicator was initially proposed by Matthews (1975) and recently regained the attention of the deep learning community. MCC is an indicator that produces a high score only if the prediction obtained good results in all aspects of correctness (i.e., true positive, true negative, false positive, false negative) while taking into account the relative proportion of categories, so that it is less sensitive to class imbalance. Therefore, we think MCC is a more honest indicator for evaluating model performance under class imbalance than Accuracy. When the model is perfect, the value of MCC is 1, indicating a perfect positive correlation between predictions and observations. With a value of 0, the model is no better than random. When the model is worse than a random model, MCC shows a negative value.

Model performance was tested on data that was simply split into a train and test part according to a ratio of 70% for training data and 30% for test data. In addition to the model performance assessment, we further tested the robustness of the model performance using 5-fold cross-validation. The data was split into five parts of equal size. In five iterations, in alternating order, one such part was denoted as test data while the others accumulate to the training data. Although typical image classification studies do not employ cross-validation, we considered this technique important for small datasets to quantitatively assess the stability of model estimates influenced by the number of samples.

### Opening the black-box: Model explanation

Explaining why a black box model made a prediction is important to understand how a prediction is made (Rudin, 2019 and Ryo et al., 2021). An untrustworthy model could make a correct prediction based on an inappropriate reason: For instance, in this study, it is possible that the model performs well, but what it learned is not a fungal colony-level trait but the handwritten time stamp on the Petri dish. One way to explain deep learning models is to use model agnostic explanations such as local interpretable model-agnostic explanations (LIME) (Ribeiro et al., 2016a). LIME is a post-hoc local surrogate model that is interpretable and can explain individual predictions.

We applied LIME for some samples that revealed a high score to inspect what the models learned visually. We set the following hyperparameters: kernel size = 6, max distance = 50, ratio = 0.5, neighborhood size = 1000, and selected features = 100. We applied a standard technique for LIME, namely, superpixel (Neubert 2015), with the quick-shift algorithm (Vedaldi and Soatto 2008).

### Settings

All models were built in python 3.6.9 using Apache’s MXNET gluon framework with GPU support CUDA-10.1. All deep learning computations were run on google colab. LIME 0.2.0.1 was run locally on a workstation in an Ubuntu Bionic Beaver environment. Each model was trained in 20 epochs of fine-tuning on the three datasets.

## Results

At the training phase, all model and dataset combinations reached a stabilized performance score (both Accuracy and MCC) after at most 20 epochs of parameter-tuning, i.e. successive model training over 20 times using the whole dataset, indicating that the duration of the training period was satisfactory (Fig. 4 as a representative case; see Fig. S1, S2, S3 for the others). At the testing phase, all model and dataset combinations showed a substantial drop in performance (both Accuracy and MCC), meaning that the models overfitted and learned false data characteristics, which are irrelevant for taxonomic group prediction.

**Figure 4.**
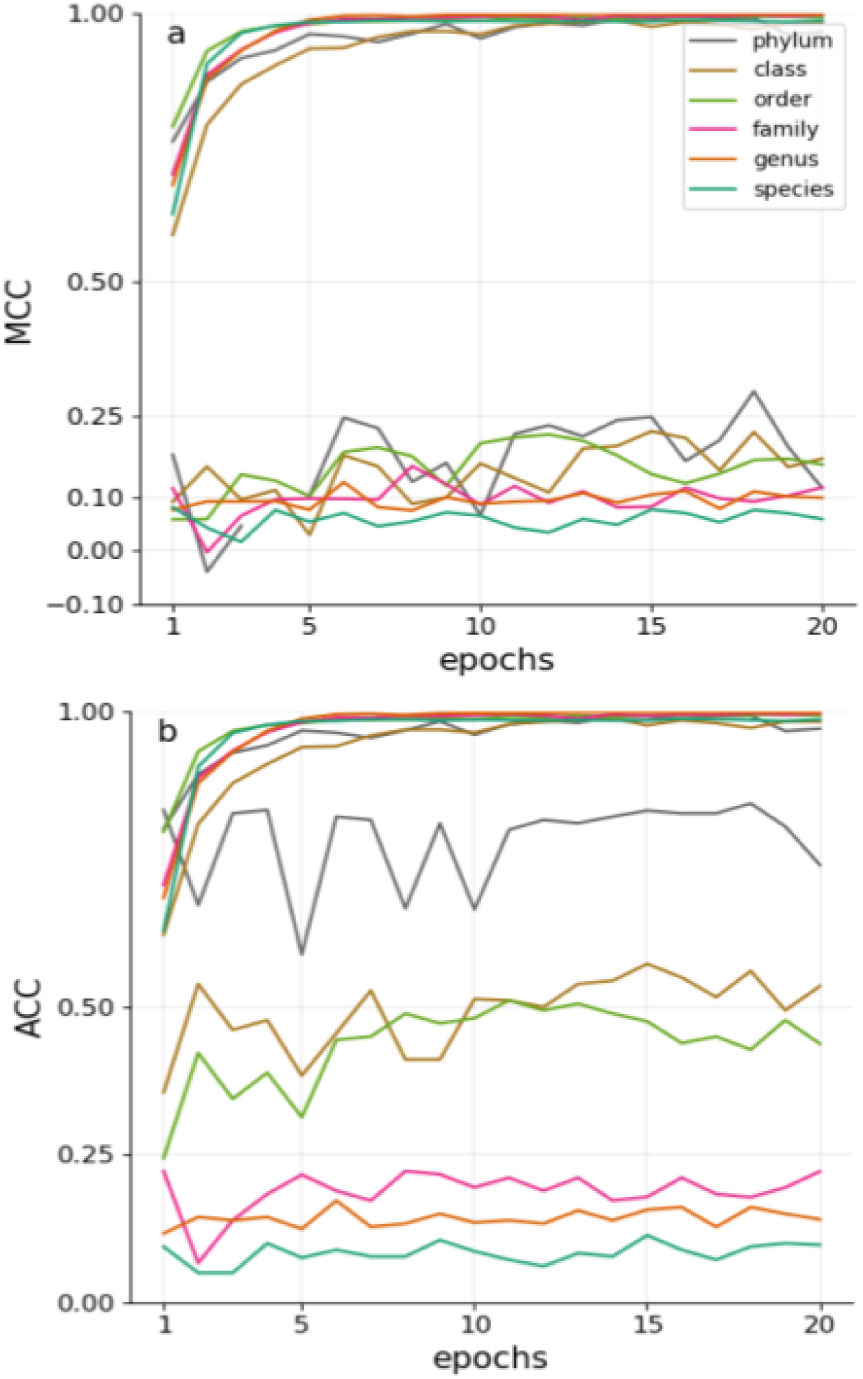
Performance for Separate Local per-level classifiers (SL) finetuned in 20 epochs on naive oversampled dataset, (a) Matthews Correlation Coefficient, (b) Accuracy. The upper curves show training scores, the lower ones are test scores.

While reducing performance in the test phase, all model-dataset combinations at the best epoch kept non-zero MCC scores, and most of them fell into the range of 0.1–0.2, meaning that the models are better than random (Fig. 5a, b). Performance scores changed along with taxonomic rank. Accuracy, as well as MCC, showed a steady decline trend from the phylum to species level. As a general trend, the SL model architecture outperformed ML and HC architectures, and HC was no better than ML. With cross-validation, all models reduced MCC scores by about 0.1 (Fig. S4, S5), but they still showed non-zero scores.

**Figure 5.**
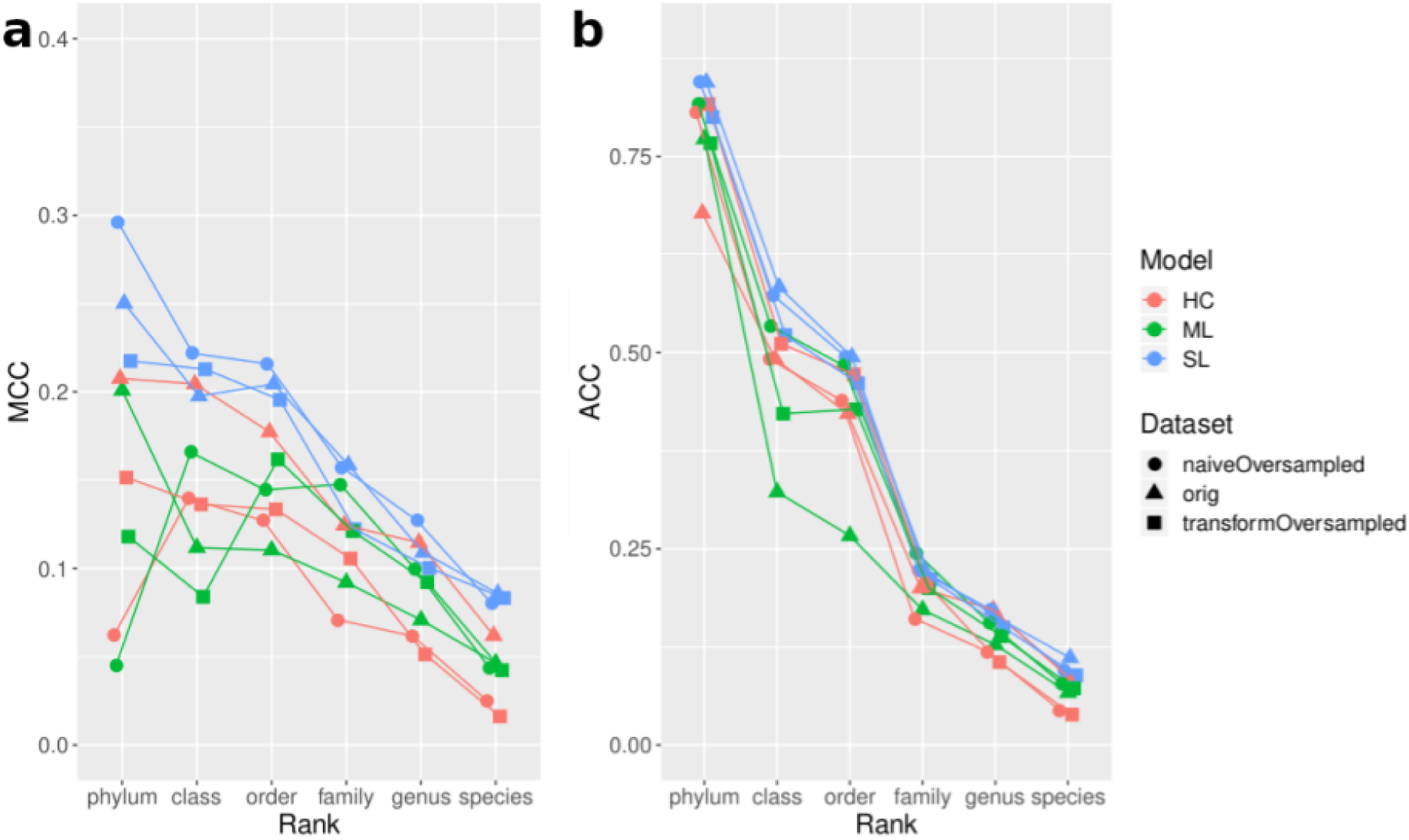
Comparison of best test performance each model achieved according to (a) Matthews Correlation Coefficient and (b) Accuracy on 606 samples.

Focusing on specific model-dataset combinations based on MCC, we observed the following. The SL with naive random oversampling performed best in three out of the six taxonomic ranks. Following the three SL models, the HC model with the original dataset was placed at the fourth performance. However, the other two HC models were no better than the ML models.

Accuracy scores reached >0.75 in many model trials at the phylum level and then linearly declined along with taxonomic resolution, reaching about 0.1. The two approaches to alleviate class imbalance, naive random oversampling and augmented random oversampling, were not substantially better than the original dataset (a difference from the original was lower than 0.05). A high Accuracy score (>0.75) seemingly suggests that the models perform very well, but indeed it mainly comes from the strong class imbalance (Fig. 3).

We found that the models learned a few taxonomic groups better than others (Fig. S6, S7, S8). At the class level, Sordariomycetes and Eurotiomycetes were well predicted, and at the order level, Hypocreales and Eurotiales were relatively well predicted. This result indicates that not all but some of fungal taxa display unique morphological characteristics at the colony level. On the other hand, this result might reflect the fact that a large majority of the isolates annotated as Hypocreales corresponded to the species *Clonostachys intermedia*, which probably exhibits a fairly consistent colony morphology.

**Figure 6.**
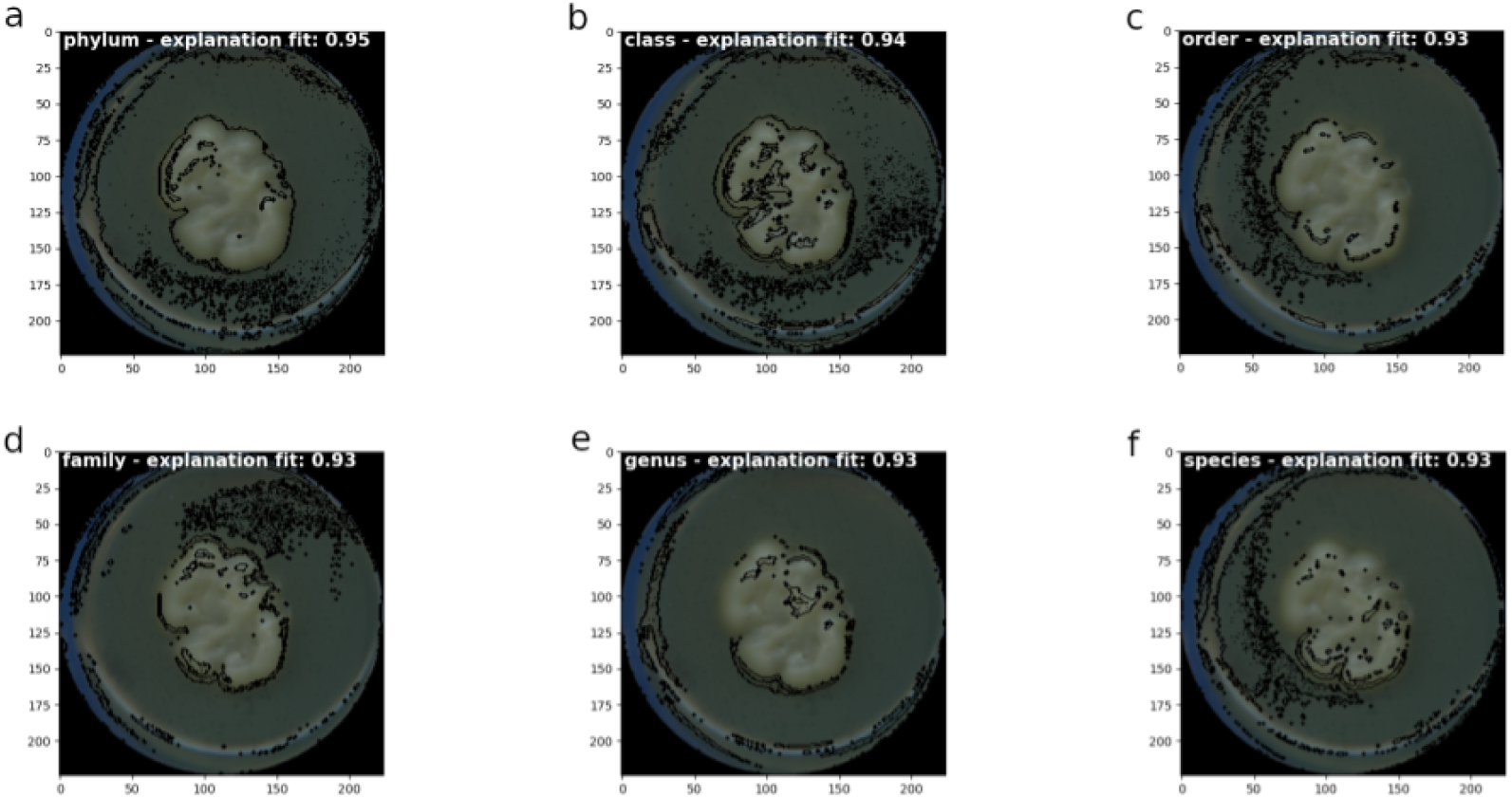
LIME explanation for *Apiotrichum dulcitum* predicted with ML at epoch 4, with neighborhood size 1000 and 100 superpixels. Segmentation is performed by quickshift algorithm with kernelsize 6, max distance 50 and ratio 0.5. Black highlighted areas are LIME explanations at rank (a) phylum, (b) class, (c) order, (d) family, (e) genus and (f) species.

To roughly grasp the explanations that led to the model’s prediction, we took some specific examples and applied LIME for images. We selected the ML model with naive random oversampling at the best performing epoch because of the intrinsic model architecture, which can learn colony traits important for classification that translate throughout taxonomic ranks. We looked at explanations for correctly classified individuals with at least five out of six taxonomic ranks. We observed that the explanation highlights a complete outline for phylum and class. Given a Petri dish surface cover after a specific time of fungal growth, a full outline could be interpreted as hyphal growth speed (Fig. 6). In lower taxonomic ranks, the outline thinned. Moreover, specific structures in the fungal colony surface were highlighted. We observed that the explanation of fungi with more complex surfaces is more challenging to interpret (Fig.S8). However, explanations seem to cluster around colony structures, too.

## Discussion

In this work, we explored the application of deep learning on taxonomically labeled image data of colonies of filamentous fungi and explanations behind the classification. We demonstrated that deep learning could classify some isolates at least at the phylum, class, and order levels, regardless of the diverse nature of their appearance and the difficulties inherent to the dataset. Evidently the outlines and inner regions of fungal colonies contributed to high prediction scores, which might represent the macroscopic expression of hyphal growth and structuring. The higher prediction scores for phylum, class, and order achieved, and the visual explanations of the predictions align with phylogenetic conservation of morphological traits observed previously (Lehmann et al.2020).

### Highly Skewed Data

Tackling class imbalance, we found that all models performed best on the naive oversampled dataset regarding performance and stability. It accelerated learning and seemed to boost test scores for phylum, class, and order while also reducing the noise’s influence. Augmented oversampled data also accelerated learning, yet at a slower pace. However, the general classification score for this set on the test data dropped dramatically, which might indicate that color is an essential feature for taxonomic identification.

We also showed that Matthews correlation coefficient (MCC) is more robust than Accuracy for evaluating the performance of deep learning models (or, more generally, any multi-classification problems in ecology). As we demonstrated, Accuracy is not a plausible indicator when the data are highly imbalanced, which is a common issue in ecology (e.g., a few species dominate the majority of relative abundance in a community). Instead, MCC is a fairer, more honest indicator for assessing model performance. An alternative is Synthetic Minority Over-sampling Technique (SMOTE) ((Chawla et al., 2002)). It offers a logical way of the synthetic creation of new samples in similarity to the existing ones.

At the class level, Sordariomycetes were well predicted. This can be attributed to the large frequency at which we recovered Clonostachys intermedia (n >150) in our culture set. This means that from a total of ∼300 isolates that were classed as Hypocreales, half belong to this species only. This unique morphological feature was probably captured during model training.

### Hierarchical Modeling

Tackling the taxonomy’s hierarchical nature, we found that separate local per-level classifiers (SL) reached a slightly higher performance score than the other models. Unexpectedly, hierarchically chained (HC) classifiers achieved the worst performance. One reason could be that employing overly complex classifiers increases the chance of error propagation rather than increasing general performance. This performance can be improved by increasing the sample size. Another possible and more ecologically exciting reason is that the inclusion of the hierarchical nature does not improve because the macroscopic morphological characteristics of fungi are phylogenetically not conserved along with taxonomic ranks. The fungal kingdom is full of examples where homologous morphological manifestations have appeared independently within disparate lineages. One example of this is the repeated transition from filamentous to yeast morphologies among distant fungal clades (Nagy et al., 2014). Another example is the existence of several pleomorphic species within the kingdom (Hibbett and Taylor 2013). It is with the wide adoption of molecular phylogenetics, and more recently phylogenomics, that fungal taxonomists have confirmed how misleading and inconsistent morphological based classifications can be, hence the calls for the modernisation of fungal systematics (Hibbett et al., 2016). On the other hand, it must be noted that fungal phylogenies are far from being clearly resolved (Naranjo-Ortiz & Gabadon, 2019). Hence the use of an imperfect hierarchical classification to train a model has to be viewed with caution. We consider, nevertheless, that ML is the most promising approach in terms of the balance between performance and possible self-learned inclusion of hierarchy.

### Small Data Set

Addressing the small sample size, we applied transfer learning and also checked performance stability with cross-validation. Cross-validation is rarely applied for deep learning approaches, but it can be done thanks to the small sample size. Given this, we could carefully check if the performance was just by chance or not. We think this is a more honest approach than showing a point estimate of a performance indicator. However, it has a clear drawback, since the sample size becomes even smaller by keeping a part of the data for validation. We conducted the same analyses after obtaining additional ca. 300 samples, but we observed no qualitative improvement from the 606 samples we investigated (see Supplementary Information Fig. S10-S13).

One possible approach to compensate for the lack of data is to add predictors that can represent some critical ecological information. In our case, for instance, technically it is possible to add the information about the period of growth time as a predictor in addition to the images, which may help the deep learning to reflect hyphal growth speed as key information that potentially improves the model performance.

### Outlook

Deep learning has proven its potential for classification analysis of image data and has found its way into ecology for different ecosystem studies and scales (Christin et al., 2018). To our knowledge, we were the first to apply an explainable artificial intelligence technique to mycelia of filamentous fungi. Improving predictive quality should be of priority for follow-up studies. This could be achieved by utilizing higher resolution images and by increasing the amount of data the algorithm can learn from while keeping class imbalance low. For example, this technique can be applied to microscopic images. Furthermore, studies that visually explore phylogenetic conservation of mycelium morphological traits can synthesize the deep learning model’s intrinsic hierarchical structure with the hierarchical structure of taxonomic data.

## Acknowledgments

We thank the managers of the three Exploratories, Kirsten Reichel-Jung, Iris Steitz, and Sandra Weithmann, Juliane Vogt, Miriam Teuscher and all former managers for their work in maintaining the plot and project infrastructure; Christiane Fischer for giving support through the central office, Andreas Ostrowski for managing the central database, and Markus Fischer, Eduard Linsenmair, Dominik Hessenmöller, Daniel Prati, Ingo Schöning, François Buscot, Ernst-Detlef Schulze, Wolfgang W. Weisser and the late Elisabeth Kalko for their role in setting up the Biodiversity Exploratories project. We thank the administration of the Hainich national park, the UNESCO Biosphere Reserve Swabian Alb and the UNESCO Biosphere Reserve Schorfheide-Chorin as well as all landowners for the excellent collaboration.The work has been partly funded by the DFG Priority Program 1374 “Biodiversity-Exploratories” (HE 6183/5-1). Field work permits were issued by the responsible state environmental offices of Baden-Württemberg, Thüringen, and Brandenburg. SH acknowledges support from Moritz Mittelbach and our student helpers Eka Giorgashvilli and Orestis Kanaris.

## Supplementary

**Figure S1.**
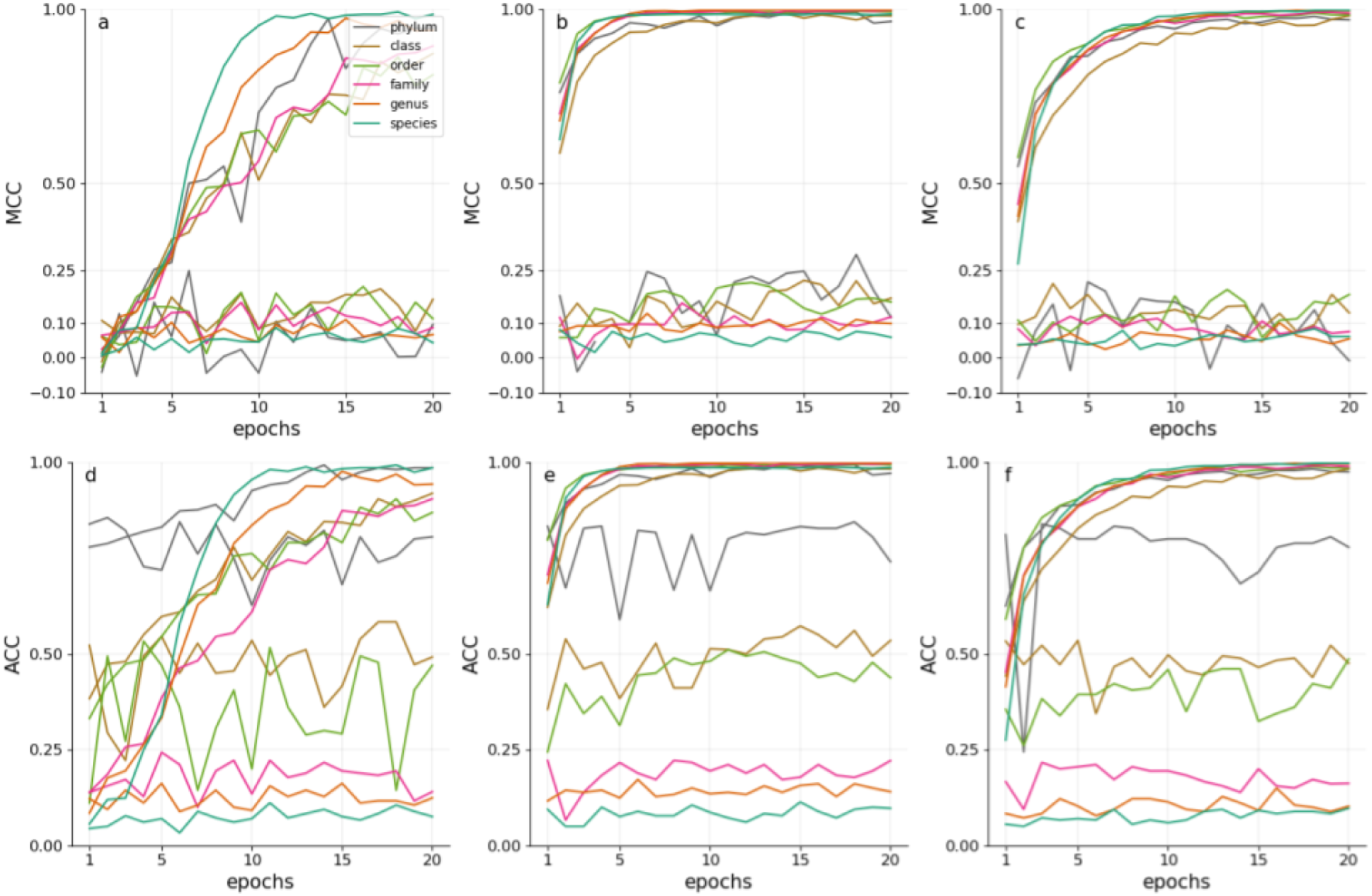
Performance for Separate Local per-level classifiers (SL) finetuned in 20 epochs according to Matthews Correlation Coefficient on (a) original, (b) naive oversampled, (c) transform oversampled data and according to Accuracy on (d) original (e) naive oversampled (f) transform oversampled data sets.

**Figure S2.**
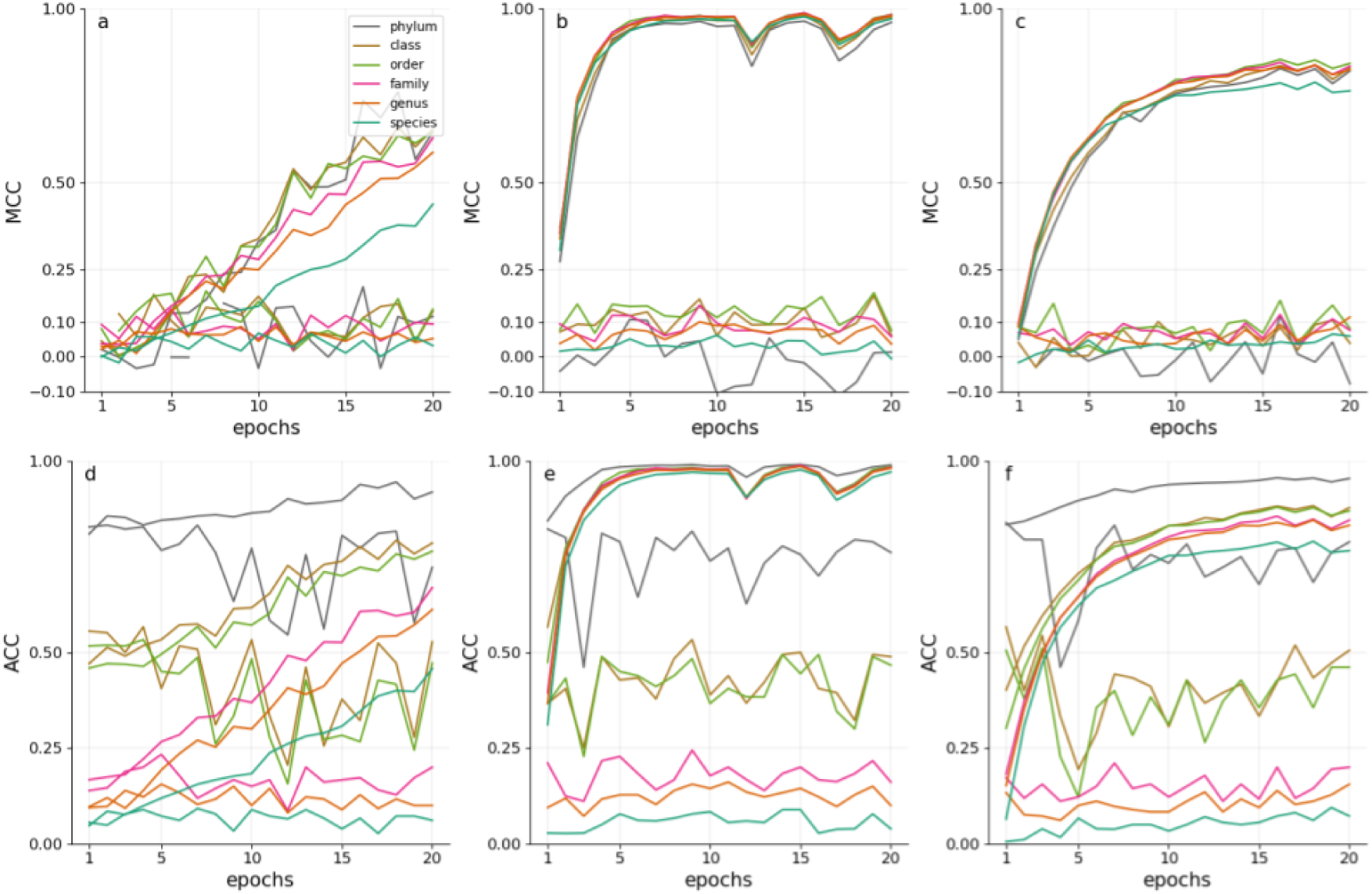
Multi-Label classifier (ML) finetuned in 20 epochs according to Matthews Correlation Coefficient on (a) original, (b) naive oversampled, (c) transform oversampled dala and according to Accuracy on (d) original (e) naive oversampled (f) transform oversampled data sets.

**Figure S3.**
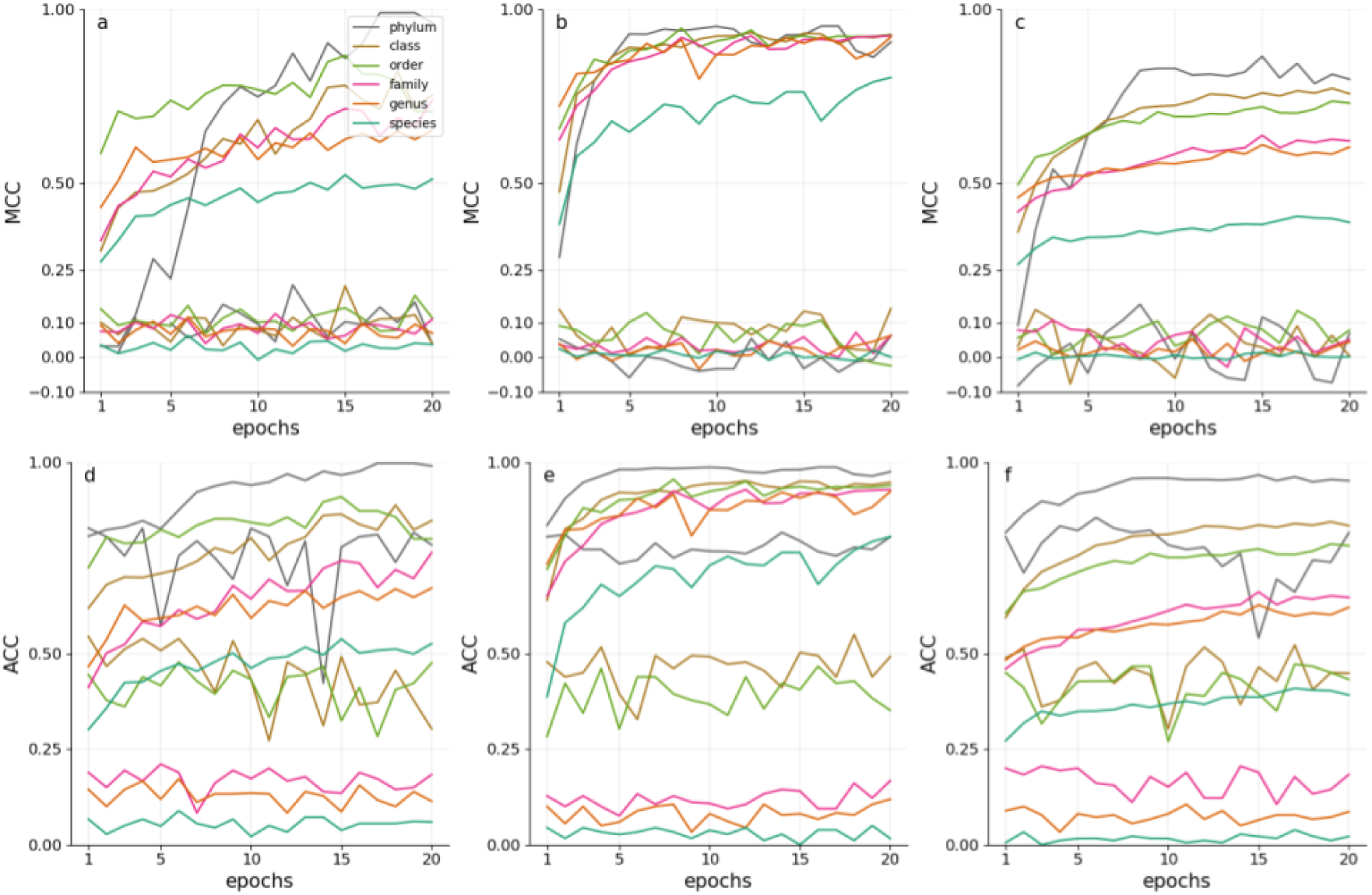
Hierarchically-Chained Local per-level classifiers (HC) finetuned in 20 epochs according to Matthews Correlation Coefficient on (a) original, (b) naive oversampled. (c) transform oversampled data and according to Accuracy on (d) original (e) naive oversampled (f) transform oversampled data sets.

**Figure S4.**
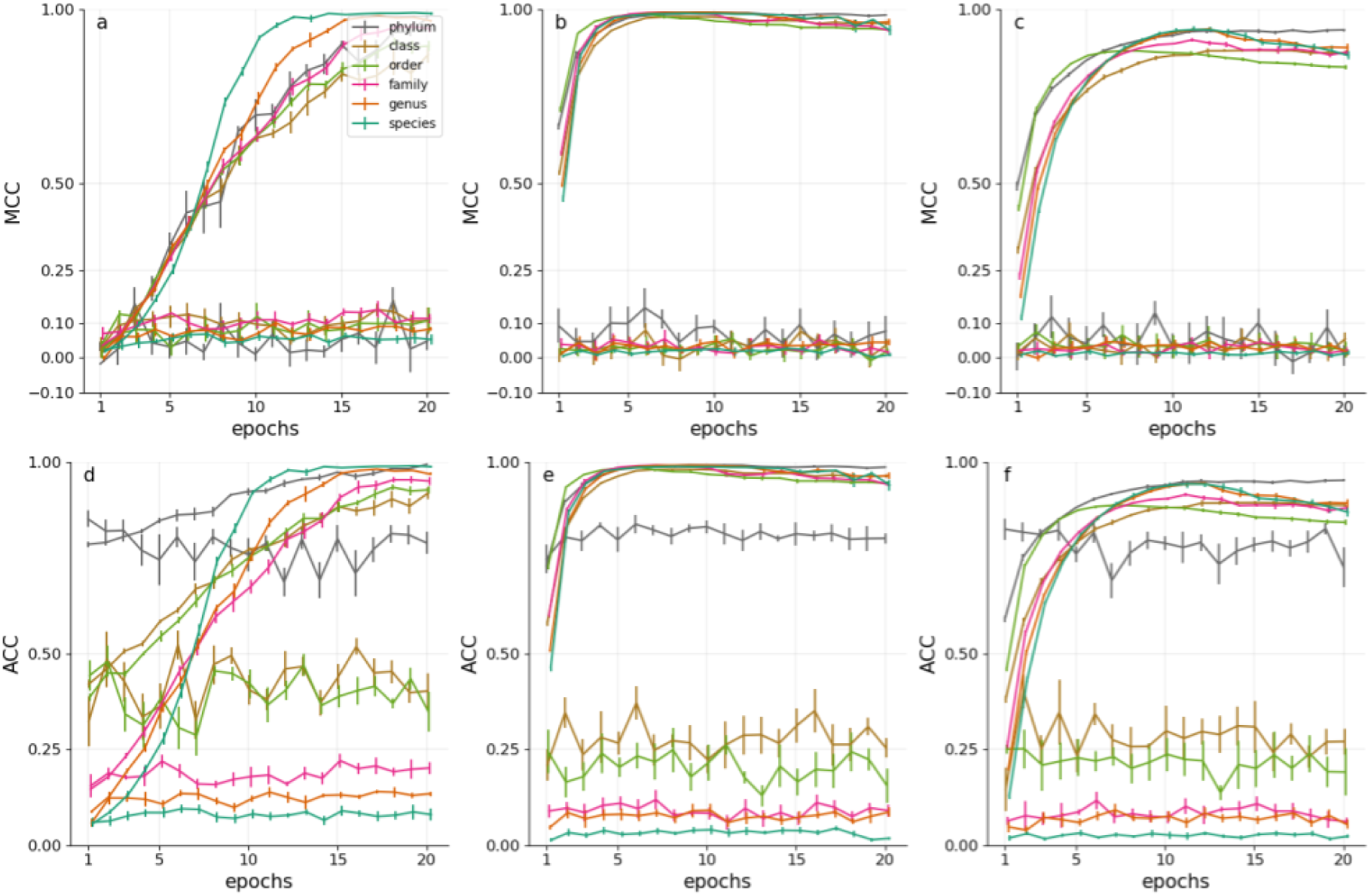
Stability for Separate Local per-level classifiers (SL) on the 606 sample set, fineluned in 20 epochs according to Matthews Correlation Coefficient on (a) original, (b) naive over-sampled, (c) transform oversampled data and according to Accuracy on (d) original (e) naive oversampled (f) transform oversampted data sets.

**Figure S5.**
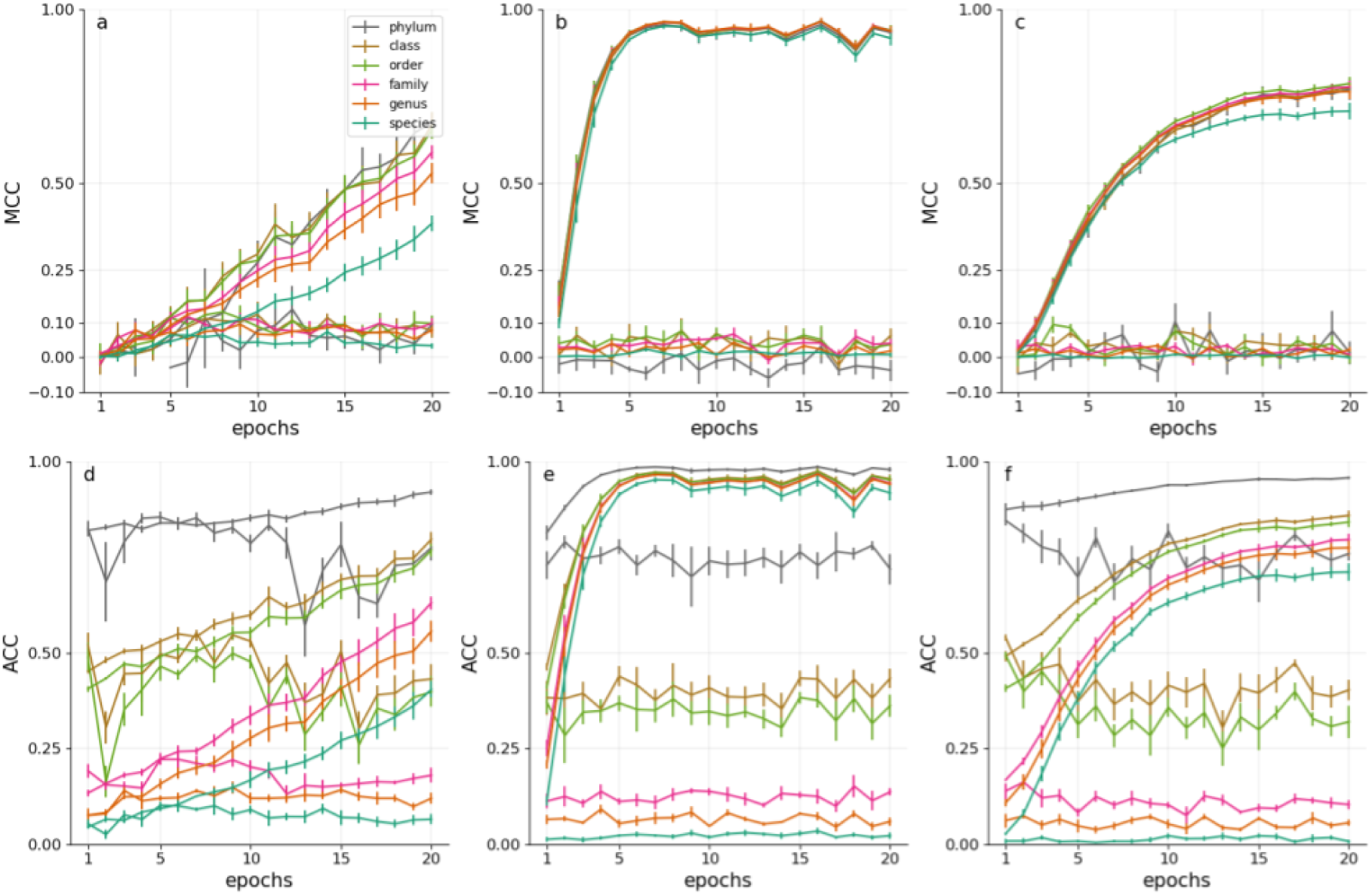
Stability for Multi Label Classifier (ML) en the 606 sample set, finetuned in 20 epochs according to Matthews Correlation Coefficient on (a) original, (b) naive oversampled, (c) transform oversampled data and according to Accuracy on (d) original (e) naive oversampled (f) transform oversampled data sets.

**Figure S6.**
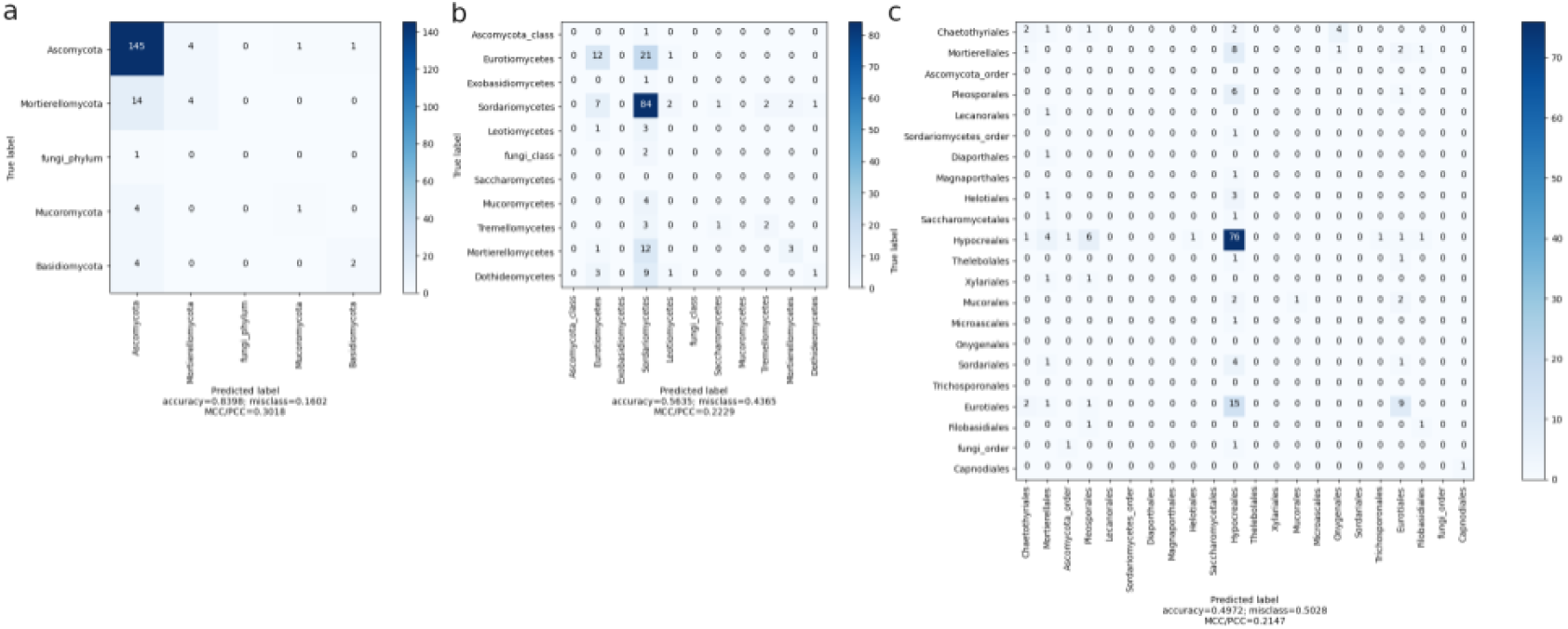
Confusion Matrices of SL model trained on the 606 sample, naive oversampled dataset Prediction on test data for taxonomic rank (a) phylum at epoch 17, (b) class at epoch 14 and (c) order at epoch 11.

**Figure S7.**
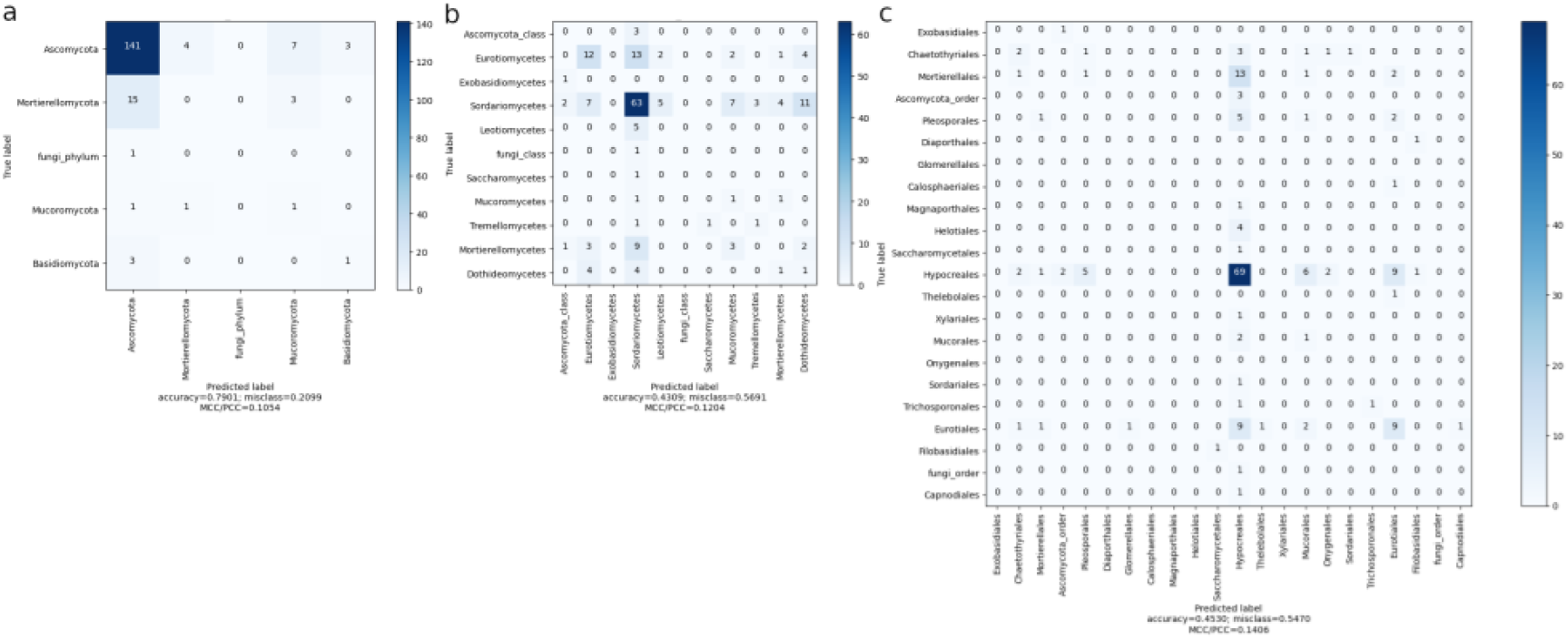
Confusion Matrices of ML model at epoch 5, trained on the 606 sample, naive oversampled dataset. Prediction on test data for taxonomic rank (a) phylum, (b) class and (c) order.

**Figure S8.**
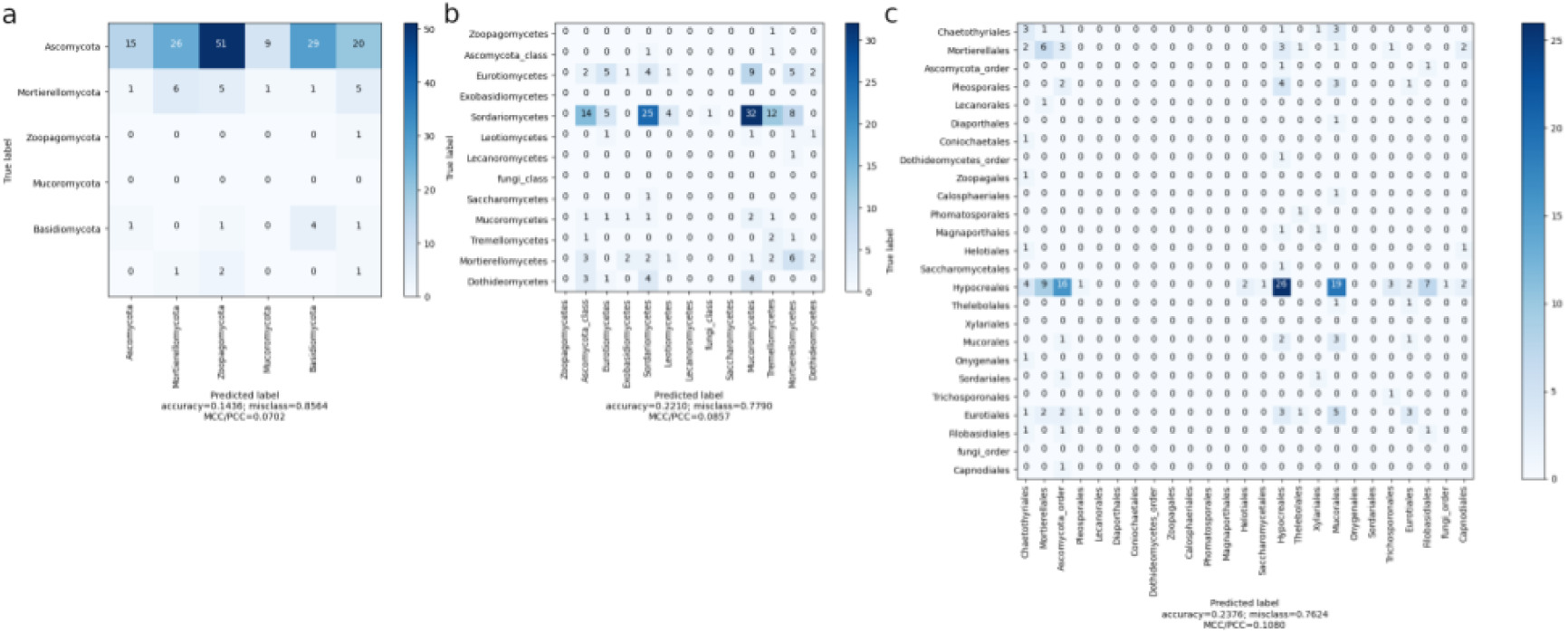
Confusion Matrices of HC model trained on the 606 sample, naive oversampled dataset. Prediction on test data for taxonomic rank (a) phylum at epoch 20. (b) class at epoch 20 and (c) order at epoch 6.

**Figure S9.**
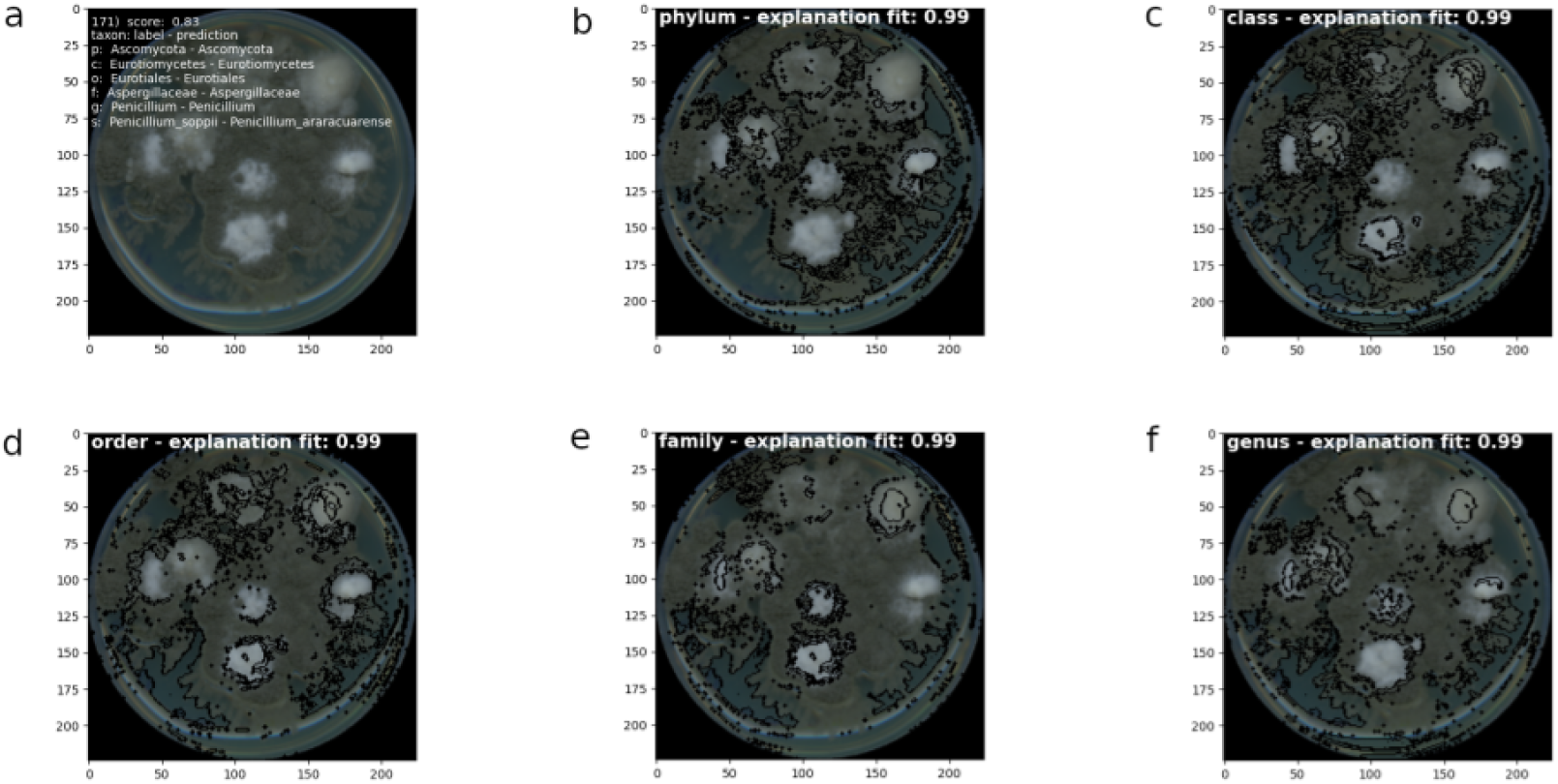
LIME explanation for *Pemcillium araracuarense* predicted with ML at epoch 4 on the 606 sample dataset, with neighborhood size 1000 and 100 superpixels. Segmentation is performed by quickshift algorithm with kernelsize 6, max distance 50 and ratio 0.5. (a) Image of the colony without explanations, listed are label and prediction with score denoting the percent of correct predictions LIME explanations are only provided for correct predictions. Black highlighted areas are LIME explanations at rank (b) phylum, (c) class, (d) order, (e) family, (f) genus.

**Figure S10.**
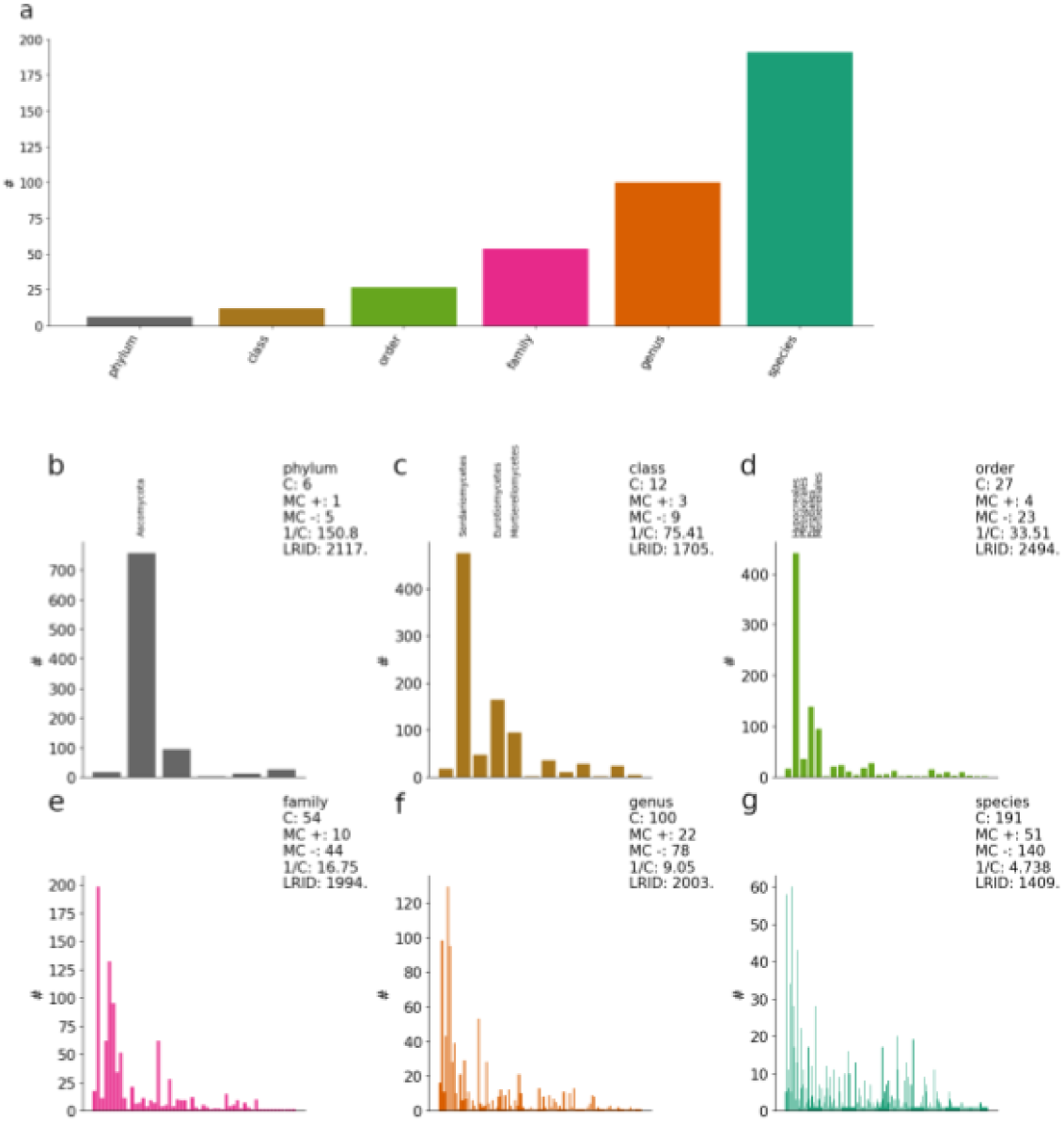
(a) Number of unique taxa/dasses according to taxonomic rank for **896 samples**. (b-g) the frequency distribution within each taxonomic rank, C is the number of categories; MC + is the majority category count, MC - is the minority category count. A category is regarded as a majority category if the number of samples # is higher than the average number of samples per category. LRID is the likelihood ratio imbalance degree For phylum, class, and order levels, the majority classes are labeled for visualization.

**Figure 11.**
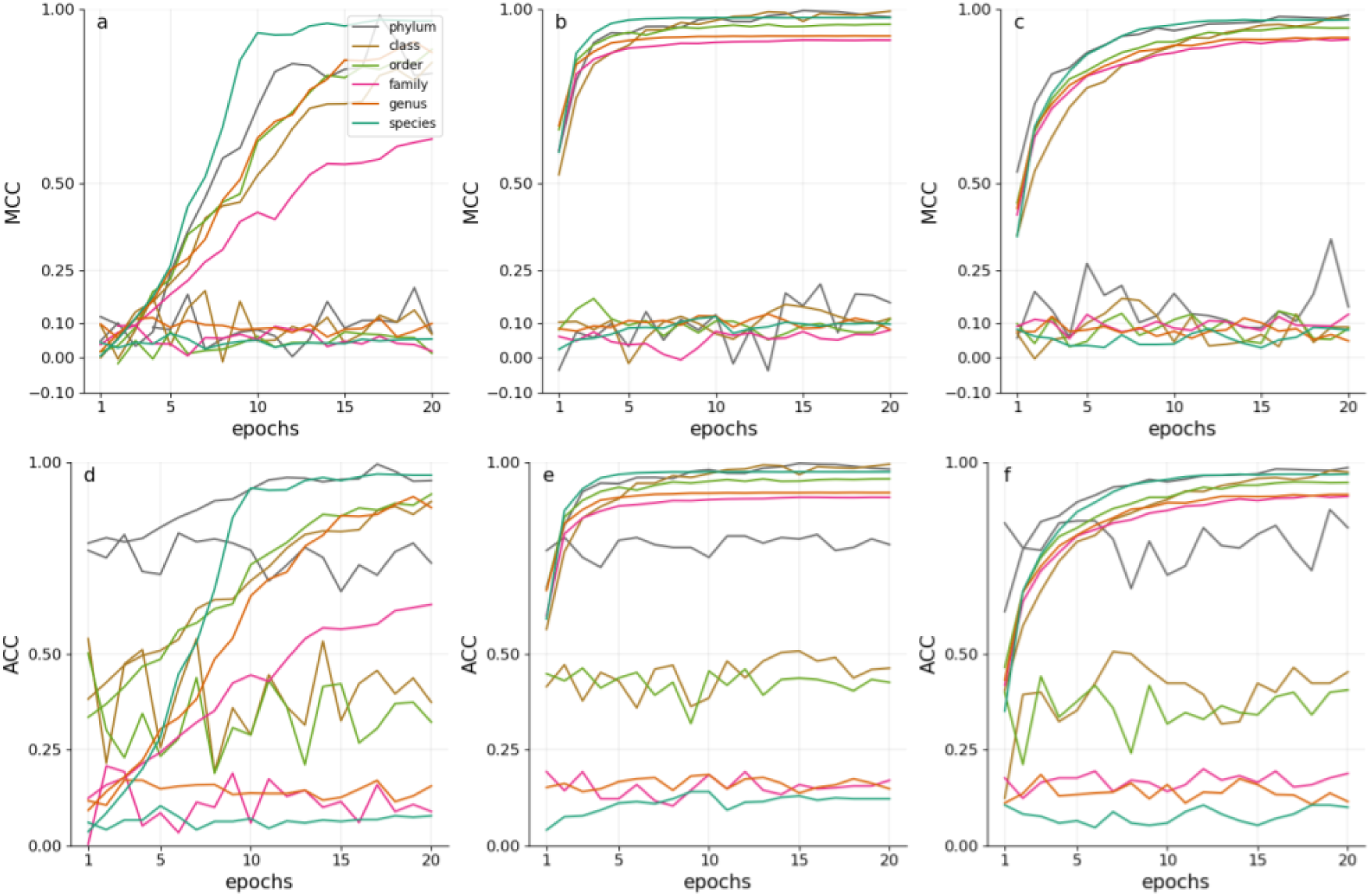
Performance for Separate Local per-level classifiers (SL) finetuned in 20 epochs on **896 samples** according to Matthews Correlation Coefficient on (a) original, (b) naive oversampled, (c) transform oversampled data and according to Accuracy on (d) original (e) naive oversampled (f) transform oversampled data sets.

**Figure S12.**
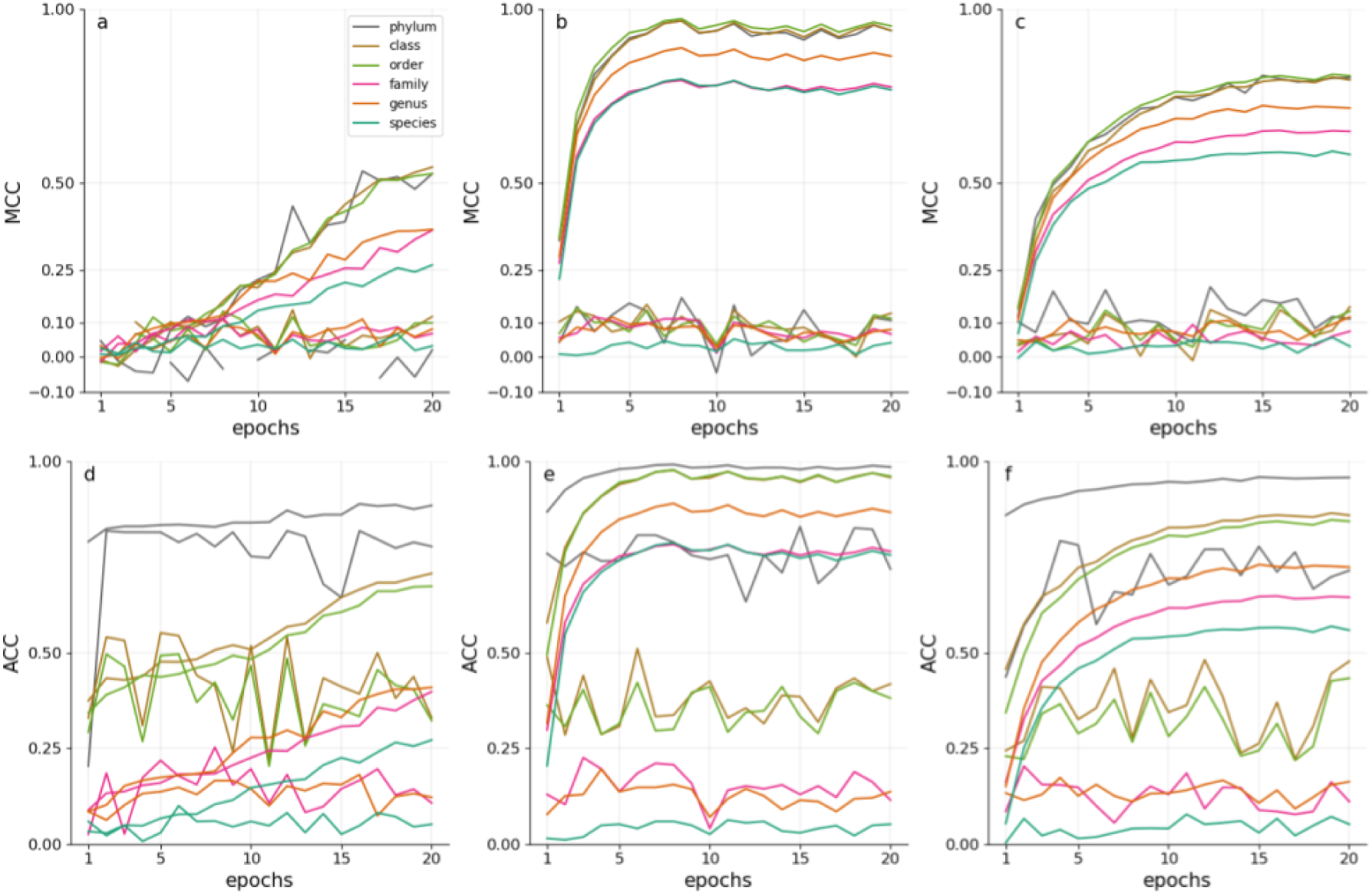
Multi-Label classifier (ML) finetuned in 20 epochs on **896 samples** according to Ma (thews Correlation Coefficient on (a) original, (b) naive oversampled. (c) transform oversampled data and according to Accuracy on id) original (e) naive oversampled (f) transform oversampled data sets.

**Figure S13.**
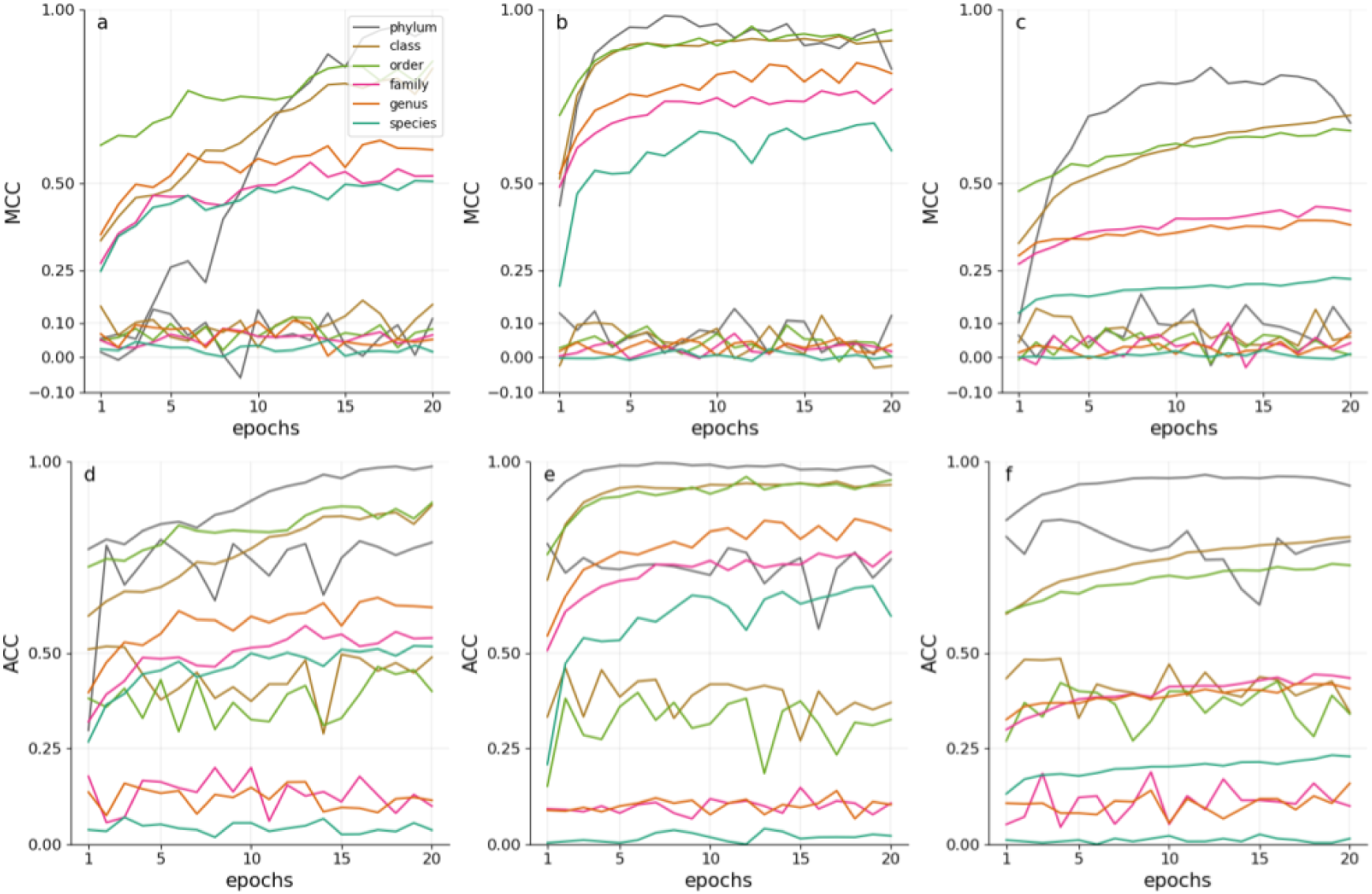
Hierarchically-Chained Local per-level classifiers (HC) finetuned in 20 epochs on **896 samples** according to Matthews Correlation Coefficient on (a) original, (b) naive oversampled, (c) transform oversampled dala and according to Accuracy on (d) original (e) naive oversampled (f) transform oversampled data sets.

